# Mucin-binding protein shuttles enable delivery of brain-targeted therapeutics

**DOI:** 10.64898/2026.03.22.713512

**Authors:** Sophia M. Shi, Gabrielle S. Tender, Jian Xiong, Josephine K. Buff, Hannah I. Park, Justin H. Mendiola, Edward N. Wilson, Monther Abu-Remaileh, Carolyn R. Bertozzi, Tony Wyss-Coray

**Affiliations:** Department of Chemistry, Stanford University, Stanford, CA, USA; Stanford Chemistry, Engineering & Medicine for Human Health (ChEM-H), Stanford University, Stanford, CA, USA; Department of Neurology and Neurological Sciences, Stanford University School of Medicine, Stanford, CA, USA; Wu Tsai Neurosciences Institute, Stanford University School of Medicine, Stanford, CA, USA; Rowland Institute, Harvard University, Cambridge, MA, USA; Department of Chemical Engineering, Stanford University, Stanford, CA, USA; Department of Genetics, Stanford University, Stanford, CA, USA; The Phil and Penny Knight Initiative for Brain Resilience, Stanford University, Stanford, CA, USA; Howard Hughes Medical Institute, Stanford University, Stanford, CA, USA

## Abstract

The blood-brain barrier (BBB) poses a major obstacle to the delivery of therapeutics into the central nervous system (CNS) due to its highly restrictive permeability. Here, we introduce glycan-targeted delivery vehicles, or GlycoShuttles, that traverse the BBB by harnessing the cerebrovascular glycocalyx, a carbohydrate-rich layer lining the BBB lumen. We discover that mucin-domain glycoproteins within this structure serve as novel entry portals for brain delivery and engineer mucin-binding protein shuttles that enable efficient transport of diverse molecular cargo across the BBB into multiple key brain cell types. This modular platform facilitates enhanced brain delivery of a variety of payloads, including antibodies and lysosomal proteins, and demonstrates therapeutic efficacy in mouse models of dementia. Our findings establish mucin-targeted GlycoShuttles as a versatile platform for noninvasive brain delivery of therapeutics, opening new avenues for the treatment of CNS diseases.

## Introduction

The blood-brain barrier (BBB) forms a highly selective vascular interface that tightly regulates the movement of molecules between the blood and brain, effectively preventing the entry of most systemically administered agents into the central nervous system (CNS). This presents a major challenge for the development of therapeutics targeting neurological and neurodegenerative diseases. Current methods that circumvent the BBB, such as intracranial, intrathecal, or intraventricular administration, are highly invasive and often result in poor biodistribution, limiting their clinical applicability (*1*). Consequently, there is a critical need for systemic delivery strategies that can enable efficient, noninvasive transport of therapeutics into the brain. Among these approaches, receptor-mediated transcytosis (RMT) has emerged as a promising strategy in clinical development, which leverages endogenous brain endothelial cell surface receptors, such as the transferrin receptor, insulin receptor, or CD98hc, that can internalize and transport ligands into the CNS without major disruption of native biological processes (*2–4*). While most brain shuttle platforms have largely focused on protein receptors as binding targets, many of these surface proteins are extensively glycosylated, and comparatively little effort has been directed toward exploiting glycan motifs themselves as targeting ligands. Notably, the luminal surface of the brain vasculature is layered in a dense, carbohydrate-rich meshwork of proteoglycans, glycoproteins, and glycolipids known as the cerebrovascular glycocalyx (*5–7*) (Fig. 1A). We hypothesized that this structure may provide an alternative set of glycan-based binding motifs capable of promoting cargo internalization into the brain.

**Figure 1.**
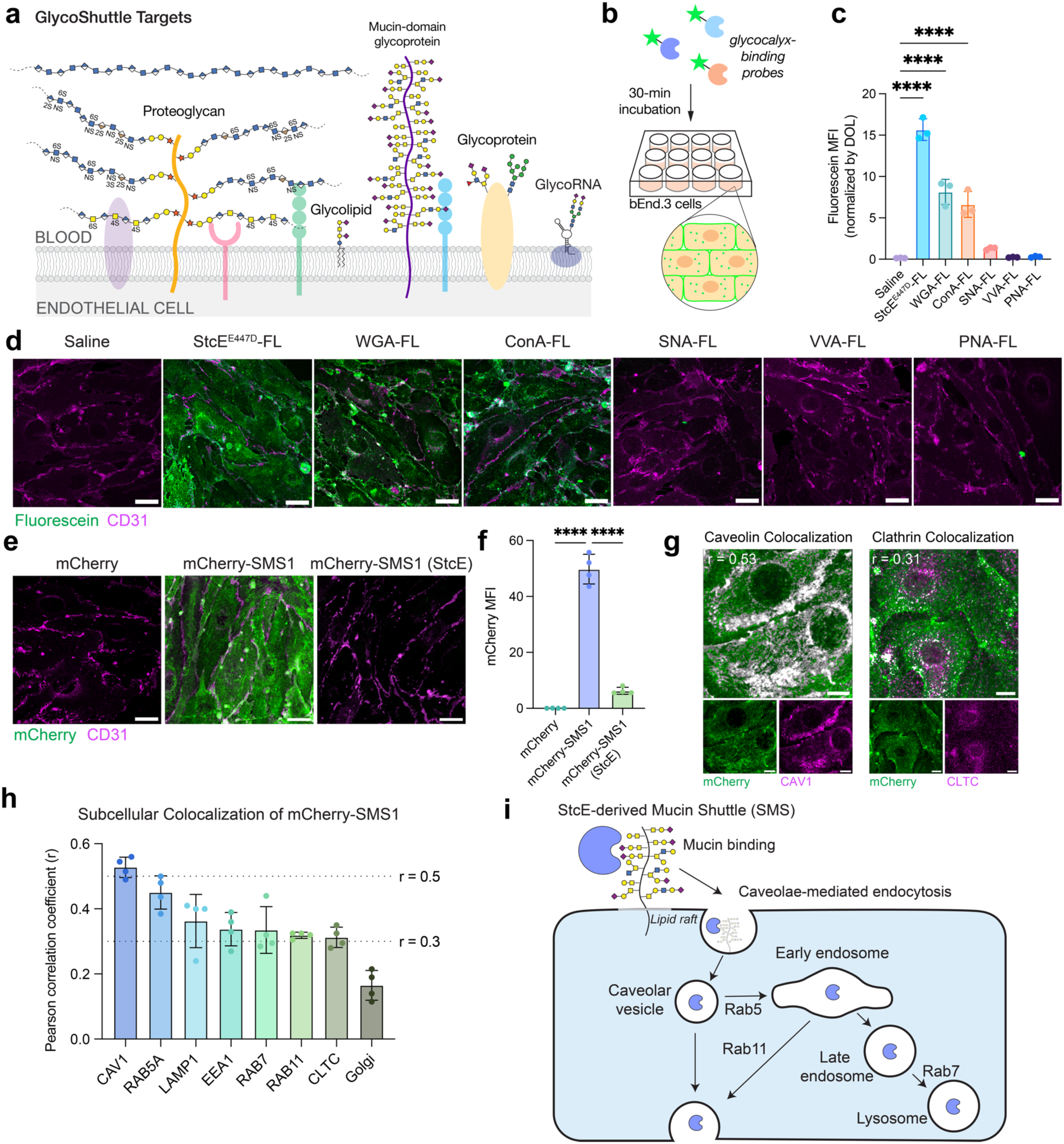
Glycocalyx-binding proteins drive cellular uptake of cargo. a) Diagram of cell surface glycocalyx components that may serve as GlycoShuttle targets. b) Cellular internalization assay to evaluate the capacity of fluorescently labeled glycocalyx-targeting probes to bind and enter bEnd.3 cells. c) Quantification of the mean fluorescence intensity (MFI) of fluorescein in cells normalized by the degree of labeling (DOL) of glycocalyx-binding probes (n=3 wells per construct; one-way ANOVA with Dunnett’s post hoc test; mean ± s.e.m.). d) Representative images of cellular binding and internalization of glycocalyx-binding probes after 30-min incubation with bEnd.3 cells. Scale bar=25 µm. e) SMS1 shuttles mCherry cargo into bEnd.3 cells in a mucin-dependent manner. Pre-treatment of cells with StcE mucinase for 1 h removes cell surface mucins (*5, 8*) and abolishes SMS1 binding and internalization. Scale bar=25 µm. f) Quantification of mCherry MFI in (e) (n=4 wells per construct; one-way ANOVA with Dunnett’s post hoc test; mean ± s.e.m.). g) mCherry-SMS1 colocalization with caveolin-1 (CAV1) and clathrin heavy chain (CLTC) in bEnd.3 cells with average Pearson correlation coefficient (r) displayed. Colocalization threshold mask in grayscale. Scale bar=10 µm. h) Colocalization of mCherry-StcE^E447D^ with various subcellular compartments measured by Pearson correlation coefficient (n=4 wells per compartment; mean ± s.e.m.). i) Schematic depicting our model of SMS internalization into brain endothelial cells. SMS binds mucins on the surface of brain endothelial cells, leading to internalization of SMS and any attached cargo in cholesterol-rich microdomains, or lipid rafts, via caveolae-mediated endocytosis. Cargo can then be trafficked via the endolysosomal pathway or transcytosed.

Despite its strategic location at the blood-brain interface and essential roles in supporting BBB function (*5*), the glycocalyx remains an underexplored target for CNS drug delivery. In this study, we demonstrate that glycocalyx-binding proteins serve as a new class of BBB-crossing shuttles, which we term GlycoShuttles, that can take advantage of the abundant glycocalyx structure on the cerebrovascular lumen to transport associated cargo across the BBB. Through this approach, we identify mucin-domain glycoproteins as novel BBB targets and utilize mucin-binding proteins to enable robust and modular brain delivery of diverse therapeutic payloads, including an anti-beta-secretase 1 (BACE1) antibody and progranulin (PGRN) lysosomal protein. These shuttles facilitate enhanced brain uptake and therapeutic efficacy in mouse models of Alzheimer’s disease (AD) and progranulin-related frontotemporal dementia (GRN-FTD), respectively. Collectively, this work establishes a distinct mucin-dependent route for BBB transport and highlights GlycoShuttles as a promising new platform for CNS drug delivery.

### Glycocalyx-binding probes undergo cellular internalization

To assess the capacity of glycocalyx-binding proteins to mediate cellular uptake in brain endothelial cells, we screened a panel of fluorescein (FL)-conjugated probes, including mucin-binding protein (StcE^E447D^) (*8*) and several plant lectins commonly used to label cell surface glycans (Fig. S1A). Following 30-minute incubations in bEnd.3 mouse brain endothelial cells, these probes displayed a broad range of internalization efficiencies and subcellular localization patterns (Fig. 1B-C). Notably, StcE^E447D^-FL, wheat germ agglutinin (WGA-FL), and concanavalin A (ConA-FL) were robustly internalized, whereas *Sambucus nigra* agglutinin (SNA-FL), *Vicia villosa* agglutinin (VVA-FL), and peanut agglutinin (PNA-FL) showed minimal to no uptake. Internalization efficiency did not correlate with the molecular weights or isoelectric points of glycocalyx-targeting probes (Fig. S1A). Furthermore, despite overlapping glycan specificities between probes (*9*), these glycocalyx-binding proteins exhibited markedly different uptake efficiencies and intracellular distributions, suggesting that additional physicochemical or structural properties influence endocytosis.

WGA and ConA have previously been reported to internalize via receptor-mediated endocytosis in various cell types, including epithelial and endothelial cells, and have been employed in drug targeting applications (*10–14*). In bEnd.3 cells, both lectins exhibit strong perinuclear punctate signal, consistent with receptor-mediated endocytosis and trafficking through the endolysosomal pathway (Fig. 1D). Interestingly, StcE^E447D^-FL also binds and internalizes efficiently into bEnd.3 cells but displays a diffuse intracellular punctate distribution, suggesting that StcE^E447D^ may engage a different internalization route or trafficking mechanism. Our group recently identified mucin-domain glycoproteins, the substrates of StcE^E447D^, as highly abundant components of the luminal brain endothelial glycocalyx (*5*), motivating us to investigate StcE^E447D^ as a potential vehicle for CNS drug delivery. Overall, these results highlight the potential of glycocalyx-binding probes to serve as shuttles for cargo transport into brain endothelial cells, the first cellular barrier to BBB penetration. Based on its uptake efficiency, distinctive trafficking profile, and lack of acute toxicity (*5, 15*), we selected StcE^E447D^ as our principal candidate for further development and refer to this engineered mucin-binding shuttle as StcE-derived Mucin Shuttle 1, or SMS1.

### SMS1 internalizes cargo into cells via caveolae and lipid rafts

To assess the potential of SMS1 to act as a cargo carrier for brain delivery, we genetically fused SMS1 to fluorescent protein mCherry (26 kDa) as a proof-of-concept payload. We found that mCherry-SMS1 was robustly internalized by both mouse and human brain endothelial cells, whereas mCherry alone showed no uptake (Fig. 1E-F and fig. S1B). mCherry-SMS1 uptake was dependent on the presence of cell surface mucins, as enzymatic removal of mucins by StcE mucinase abolished internalization. In contrast, the transferrin receptor-binding antibody (anti-TfR1 [8D3]), a well-characterized brain shuttle, retained its uptake capacity regardless of mucin removal and localized to perinuclear compartments, consistent with its known clathrin-mediated endocytosis and trafficking through the endolysosomal pathway (*16*) (Fig. S1C-D).

To elucidate the internalization mechanism of SMS1, we examined the subcellular localization of mCherry-SMS1 following 30-minute incubation with bEnd.3 cells. mCherry-SMS1 exhibited strong colocalization with caveolin-1 (CAV1), a marker of caveolae, and weaker colocalization with clathrin heavy chain, indicating preferential uptake via caveolae-mediated endocytic pathways over clathrin-dependent routes (Fig. 1G). Additionally, we observed substantial colocalization of mCherry-SMS1 with endosomal and lysosomal markers, suggesting subsequent trafficking through the endolysosomal system (Fig. 1H). Pharmacological perturbations further supported this internalization model. Treatment of bEnd.3 cells with bafilomycin, an inhibitor of endolysosomal acidification, led to widespread accumulation of mCherry puncta across cells, consistent with reduced lysosomal degradation and impaired trafficking (Fig. S1E-F). Disruption of cholesterol-rich membrane domains, or lipid rafts, with nystatin, a cholesterol-sequestering agent, caused mCherry-SMS1 to form bright cellular clusters, suggesting cholesterol-dependent membrane binding (Fig. S1G). Similarly, treatment with methyl-β-cyclodextrin (MβCD), which depletes membrane cholesterol, substantially reduced the number of mCherry-positive puncta compared to saline-treated controls, further supporting a requirement for cholesterol-rich membrane domains for SMS1-mediated internalization (Fig. S1G). Together, these findings indicate that SMS1 binds to mucin-domain proteins on the endothelial cell surface and internalizes primarily via lipid raft- and caveolae-mediated endocytosis, followed by trafficking through the endolysosomal pathway (Fig. 1I).

### SMS1 mediates cargo transport into the brain

We next assessed the ability of SMS1 to mediate transcytosis of mCherry across the BBB and enter different brain cell types *in vivo*. We administered 5 mg/kg of mCherry-SMS1 or an equimolar dose of unconjugated mCherry intravenously into mice and evaluated brain uptake following transcardial perfusion at multiple time points up to 48 hours post-injection (Fig. 2A). Flow cytometry analysis revealed that SMS1 significantly enhanced mCherry delivery into brain endothelial and parenchymal cells at both 24- and 48-hours post-injection compared to unconjugated mCherry (Fig. 2B). These findings were corroborated by immunofluorescence imaging, which showed mCherry-SMS1 displaying a time-dependent pattern of internalization, with a strong vascular signal at 2 hours post-injection, transitioning to widespread, punctate signal within the blood vessels and brain parenchyma by 6 hours, which further intensified throughout the parenchyma at 24 hours (Fig. 2C-D). In contrast, mice treated with unconjugated mCherry showed no detectable mCherry signal in the brain. These observations indicate that SMS1 facilitates progressive transcytosis and dispersion of cargo in the brain over time. Co-staining with cell type–specific markers confirmed that mCherry-SMS1 was internalized by multiple brain cell types, including microglia, neurons, and astrocytes (Fig. 2E-G).

**Figure 2.**
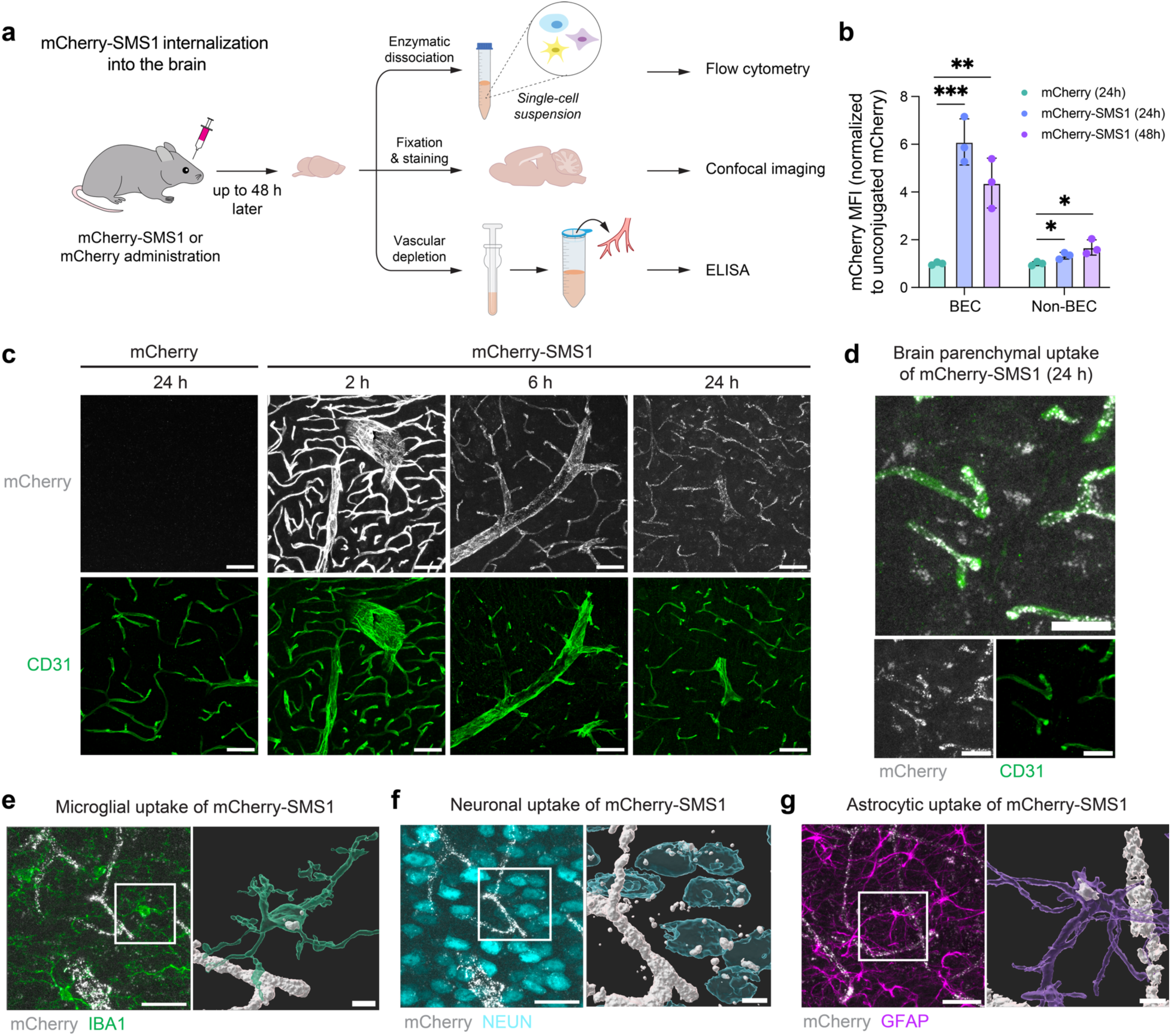
SMS1 mediates the transport of mCherry cargo into the brain. a) Experimental setup for measuring brain uptake of mCherry-SMS1 into different cell types via flow cytometry, confocal imaging, and ELISA. b) mCherry MFI in brain endothelial cell (BEC) (CD31^+^) and non-BEC (CD31^-^) populations as measured by flow cytometry at 24 h and 48 h after 5 mg/kg i.v. injection of mCherry-SMS1 or an equimolar dose of unconjugated mCherry (n=3 mice per group; one-way ANOVA with Dunnett’s post hoc test; mean ± s.e.m.). c) mCherry-SMS1 internalization into the cortex at 2, 6, and 24 h after 5 mg/kg i.v. injection compared to unconjugated mCherry internalization at 24 h. Scale bar=50 µm. d) mCherry-SMS1 internalization outside of the vasculature at 24 h after 5 mg/kg i.v. injection. Scale bar=10 µm. e) Microglial uptake of mCherry-SMS1. Scale bar=25 µm (left) and 5 µm (right). f) Neuronal uptake of mCherry-SMS1. Scale bar=25 µm (left) and 5 µm (right). g) Astrocytic uptake of mCherry-SMS1. Scale bar=25 µm (left) and 5 µm (right).

### The C-terminal domain mediates SMS1 internalization

To improve the utility of SMS1 as a brain shuttle, we sought to engineer smaller domain variants of SMS1 that retained effective trafficking properties. SMS1 is an enzymatically inactive mucinase derived from *E. coli* that exhibits high binding specificity for mucins (*17, 18*). The full-length protein comprises three globular domains and has a molecular weight of ∼98 kDa, which is relatively large for a protein shuttle. Protein shuttles of larger molecular size can limit cargo capacity, increase immunogenicity risk, and impact biodistribution. To identify the minimal functional unit required for cellular uptake, we generated SMS1 deletion constructs lacking either the C-terminal domain (SMS1ΔC) or the intervening INS region (SMS1ΔINS), both of which have been previously implicated in substrate recognition and binding (*17, 19*) (Fig. 3A-B and fig. S2A). In internalization assays using bEnd.3 cells, SMS1ΔINS maintained robust uptake, whereas SMS1ΔC exhibited a complete loss of binding and internalization, indicating that the C-terminal domain is essential for these functions (Fig. 3C-D). To test whether the C-terminal domain alone was sufficient for delivery, we generated an mCherry fusion to the isolated C-terminal domain, termed mCherry-SMS2. Remarkably, mCherry-SMS2 demonstrated efficient internalization into both mouse and human brain endothelial cells, comparable to full-length SMS1 (Fig. 3C-D and fig. S2B-C).

**Figure 3.**
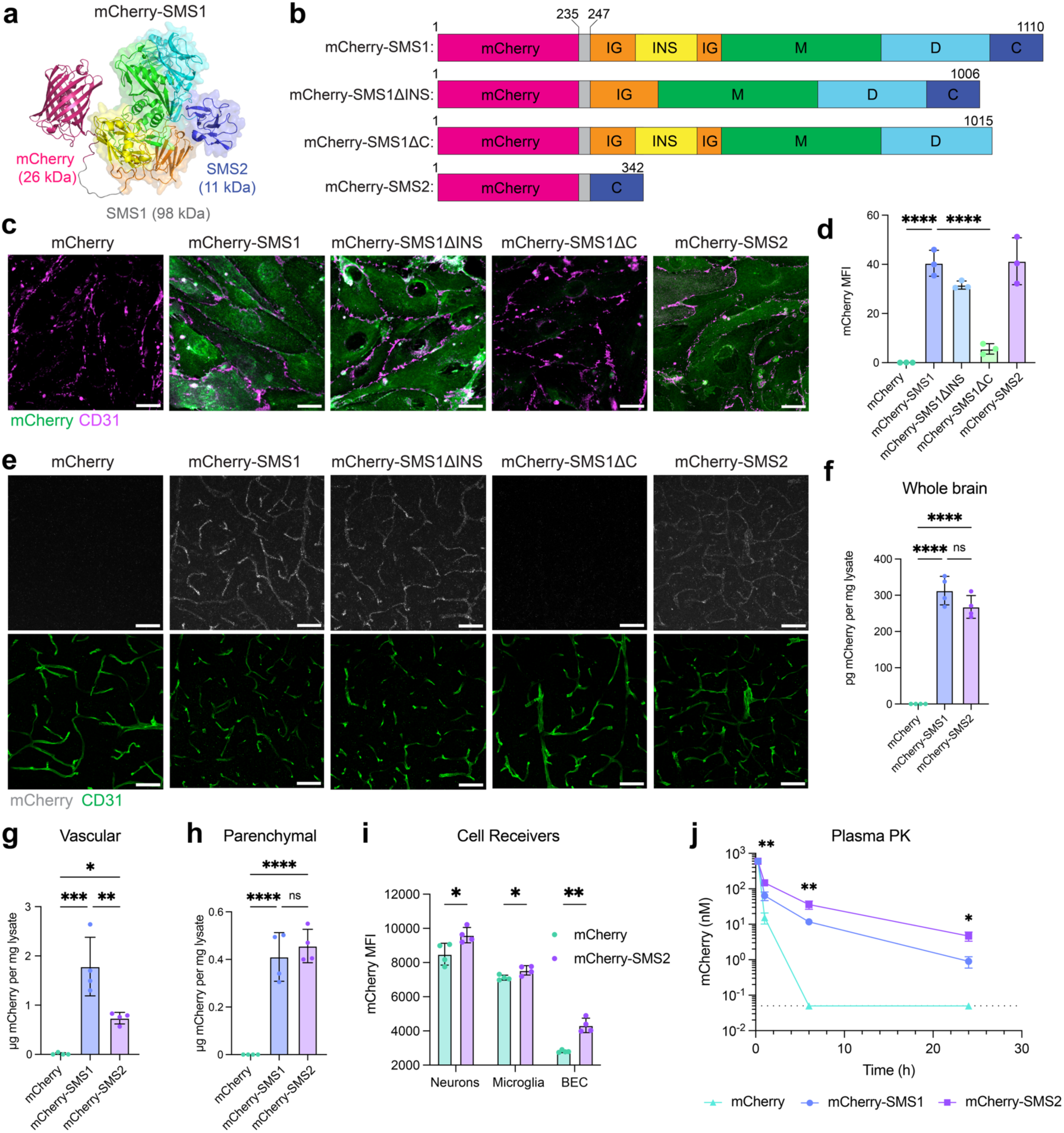
The C-terminal domain of SMS1 is necessary and sufficient for brain uptake. a) AlphaFold generated structure of mCherry-SMS1. b) mCherry-SMS1 mutant constructs used in internalization assays. Domain boundaries and nomenclature follow Yu A.C.Y, et al. (*17*). Color-coded regions include: mCherry (pink), flexible linker (gray), immunoglobulin-like domain (IG; orange), insertion region (INS; yellow), metalloprotease domain (M; green), additional disordered regions (D; cyan), and C-terminal domain (C; blue). The N-terminal His tag is omitted. Amino acid positions are indicated. c) Binding and internalization of mCherry-SMS1, mCherry-SMS1ΔINS, mCherry-SMS1ΔC, mCherry-SMS2, and mCherry in bEnd.3 cells after 30-min incubation at 37°C. Scale bar=25 µm. d) Quantification of (c) (n=3 wells per construct; one-way ANOVA with Dunnett’s post hoc test; mean ± s.e.m.). e) Internalization of mCherry-SMS1, mCherry-SMS1ΔINS, mCherry-SMS1ΔC, mCherry-SMS2, and mCherry into the cortex at 24 h after i.v. injection of 5 mg/kg mCherry-SMS1 or molar equivalents of mutant constructs. f) ELISA quantification of mCherry in brain hemisphere tissue from mice treated with mCherry, mCherry-SMS1, or mCherry-SMS2 (n=4 animals per construct; one-way ANOVA with Tukey’s post hoc test; mean ± s.e.m.). g) ELISA quantification of mCherry in the vascular fraction of brain tissue from mice treated with mCherry, mCherry-SMS1, or mCherry-SMS2 (n=4 animals per construct; one-way ANOVA with Tukey’s post hoc test; mean ± s.e.m.). h) ELISA quantification of mCherry in the parenchymal fraction of brain tissue from mice treated with mCherry, mCherry-SMS1, or mCherry-SMS2 (n=4 animals per construct; one-way ANOVA with Tukey’s post hoc test; mean ± s.e.m.). i) Flow cytometry analysis of mCherry-SMS2 uptake by neurons, microglia, and brain endothelial cells compared to unconjugated mCherry (n=4 animals per construct; two-sided t-test; mean ± s.e.m.). j) Plasma PK analysis of mCherry, mCherry-SMS1, and mCherry-SMS2-injected mice over 24 hours (n=4 animals per construct; two-way ANOVA with Dunnett’s post hoc test; mean ± s.e.m.).

Binding affinity assays in K562 cells demonstrated similar affinity of mCherry-SMS1 and mCherry-SMS2 in the middle nanomolar range (K_d_ = 364 and 265 nM, respectively) (Fig. S2D). Importantly, binding of both SMS constructs was mucin-dependent, as enzymatic removal of mucins significantly reduced mCherry signal. Mechanistically, mCherry-SMS2 colocalizes more strongly with CAV1 compared to CLTC, and mCherry-SMS2 punctate signal was significantly reduced in CAV1-knockout (KO) bEnd.3 cells, supporting a caveolae-mediated cellular entry mechanism (Fig. S2E–H). These findings establish SMS2, at ∼11 kDa, as a compact and efficient mucin-binding module sufficient to recapitulate the cellular uptake properties of full-length SMS1.

### SMS2 functions as an effective brain shuttle *in vivo*

We next compared the *in vivo* brain delivery efficiency of SMS1 domain variants following intravenous injection in mice. At 24 hours post-injection, mCherry-SMS1, mCherry-SMS1ΔINS, and mCherry-SMS2 all demonstrated robust brain uptake, whereas mCherry-SMS1ΔC and unconjugated mCherry showed no detectable signal, mirroring the observations in brain endothelial cells *in vitro* (Fig. 3E). Quantification of mCherry levels in brain hemispheres by ELISA revealed similar levels of brain uptake between SMS1 and SMS2 constructs and negligible uptake of unconjugated mCherry (Fig. 3F). To quantify vascular versus parenchymal distribution of SMS constructs, we used vascular depletion of brain tissues and confirmed effective depletion of brain endothelial cell marker CD31 from parenchymal fractions (Fig. S3A). mCherry-SMS2 showed less vascular retention compared to mCherry-SMS1 but similar levels of brain parenchymal delivery at 48 hours post-injection (Fig. 3G-H). Flow cytometry analysis of dissociated brain tissue confirmed cellular uptake of mCherry-SMS2 into key brain cell types, including brain endothelial cells, neurons, and microglia, supporting its potential for therapeutic delivery applications (Fig. 3I).

We additionally examined the distribution of SMS1 and SMS2 constructs in peripheral organs and plasma. mCherry-SMS1 displayed strong signal in liver hepatocytes and Kupffer cells, whereas mCherry-SMS2 exhibited more moderate liver uptake over unconjugated mCherry. Both SMS constructs exhibited minimal parenchymal uptake in the heart and kidney compared to unconjugated mCherry (Fig. S4A-F). Overall, SMS2 exhibited lower accumulation in the selected peripheral organs than SMS1, suggesting a more favorable brain-to-periphery delivery profile. Additionally, plasma pharmacokinetic (PK) analysis revealed extended circulation half-lives for both mCherry-SMS1 (t_1/2_ = 4.4 h) and mCherry-SMS2 (t_1/2_ = 4.8 h) compared to unconjugated mCherry (t_1/2_ = 1.0 h) (Fig. 3J). Given that mCherry-SMS2 is approximately one-third of the size of mCherry-SMS1, the longer half-life of mCherry-SMS2 may reflect its higher binding affinity to cell surfaces which may potentially prolong systemic retention. Together, these results demonstrate that SMS2 retains the brain delivery efficiency of full-length SMS1, while exhibiting reduced peripheral tissue uptake. Its compact size, broad brain distribution, uptake into major brain cell types, and favorable pharmacokinetic profile establish SMS2 as a promising and efficient platform for targeted CNS drug delivery.

### Anti-BACE1 antibody delivery and therapeutic efficacy in AD mouse model

Antibodies are attractive therapeutic agents due to their high specificity, tunable structure, and extended half-life, however, their clinical application for CNS diseases is severely limited by their large size and poor permeability across the BBB (*20, 21*). To assess the ability of SMS2 to effectively deliver antibody cargo into the brain with therapeutic activity, we engineered SMS2-fused versions of a human monoclonal antibody targeting BACE1, a transmembrane protease involved in amyloid beta (Aβ) generation and a well-established therapeutic target in AD (*3, 22, 23*). Although BACE1 inhibitors have underperformed clinically, insufficient BBB penetration remains a potential contributing factor to their limited efficacy. Importantly, for brain shuttle evaluation, BACE1 targeting offers the advantage of clear pharmacodynamic (PD) biomarkers, such as brain Aβ peptide levels, to assess target engagement.

We generated SMS2 fusions to the C-terminus of the human anti-BACE1 antibody (αBACE1) and tested internalization efficacy in bEnd.3 cells (Fig. 4A). Unconjugated αBACE1 showed no detectable binding or internalization, consistent with its limited ability to engage and internalize into brain endothelial cells (Fig. 4B-C). However, αBACE1-SMS2 exhibited significantly higher levels of binding and internalization and demonstrated a requirement for cell surface mucins for engagement. We next evaluated brain delivery *in vivo* by injecting mice intravenously with αBACE1 or αBACE1-SMS2 and assessed human IgG (hIgG) signal in brain slices by immunofluorescence staining. As expected, αBACE1 alone resulted in minimal brain uptake, with faint signal restricted to periventricular and meningeal regions (Fig. 4D). In contrast, αBACE1-SMS2 showed widespread uptake across brain regions, indicating robust BBB transport and parenchymal distribution.

**Figure 4.**
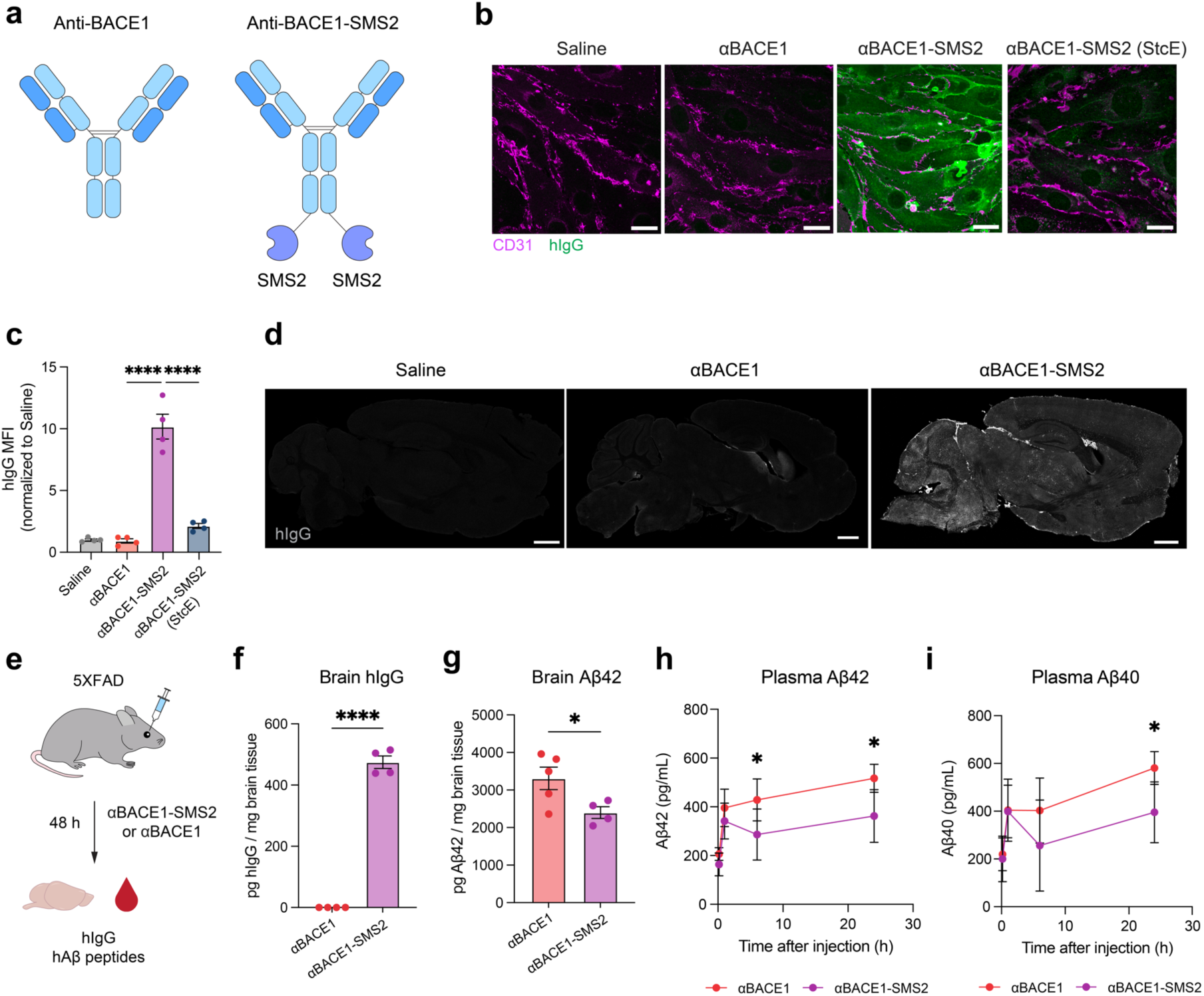
SMS2 delivers anti-BACE1 antibody into the brain and reduces amyloidogenic Aβ production. a) αBACE1 and αBACE1-SMS2 constructs used in this study. b) Binding and internalization of αBACE1 and αBACE1-SMS2 in bEnd.3 cells. Scale bar=25 µm. c) Quantification of binding and internalization of αBACE1 and αBACE1-SMS2 in bEnd.3 cells (n=4 wells per construct; one-way ANOVA with Šidák’s post hoc test; mean ± s.e.m.). Binding and internalization of αBACE1-SMS2 is dependent on cell surface mucins. d) hIgG immunofluorescence staining of mouse brains 48 hours after a single intravenous dose of saline, αBACE1 antibody, or αBACE1-SMS2 at equimolar dosing (equivalent to 25 mg/kg αBACE1-SMS2). Scale bar=1 mm. e) Experimental scheme comparing internalization and therapeutic efficacy of αBACE1 versus αBACE1-SMS2 treatment in 5XFAD mice. f) ELISA quantification of hIgG internalization in the brains of 5XFAD mice treated with αBACE1 or αBACE1-SMS2 48 hours after treatment (n=4-5 animals per construct; two-sided t-test; mean ± s.e.m.). g) ELISA quantification of Aβ42 levels in the brains of 5XFAD mice treated with αBACE1 or αBACE1-SMS2 48 hours after treatment (n=4-5 animals per construct; two-sided t-test; mean ± s.e.m.). h) Plasma PK analysis of Aβ42 levels in mice treated with αBACE1 or αBACE1-SMS2 over 24 hours (n=4 animals per construct; two-way ANOVA with Dunnett’s post hoc test; mean ± s.e.m.). i) Plasma PK analysis of Aβ40 levels in mice treated with αBACE1 or αBACE1-SMS2 over 24 hours (n=4 animals per construct; two-way ANOVA with Dunnett’s post hoc test; mean ± s.e.m.).

We next assessed therapeutic efficacy in the 5XFAD mouse model of AD, which exhibits early, aggressive amyloid pathology driven by transgenic overexpression of human amyloid precursor protein (APP) and presenilin 1 (PS1) with five familial AD mutations under the mThy1 promoter (*24*). In this model, BACE1 initiates amyloidogenic Aβ production by cleaving APP into sAPPβ and membrane-bound C99, which is subsequently processed by γ-secretase into Aβ peptides with a strong bias toward the pathogenic Aβ42 isoform. Using ELISA quantification of hIgG in brain lysates, we demonstrate that SMS2 effectively shuttles αBACE1 into the brains of 5XFAD mice, with approximately 475 pg of hIgG per mg of brain tissue detected 48 hours after 25 mg/kg αBACE1-SMS2 injection, whereas unconjugated αBACE1 was undetectable (Fig. 4E-F). Importantly, treatment with αBACE1-SMS2 led to a significant reduction in brain Aβ42 levels at 48 hours post-injection relative to αBACE1 alone, indicating effective BACE1 inhibition *in vivo* (Fig. 4G). Additionally, plasma Aβ42 and Aβ40 levels were reduced up to 24 hours post-injection in αBACE1-SMS2-treated mice compared to unconjugated αBACE1-treated mice, further supporting enhanced target engagement (Fig. 4H-I). These findings establish SMS2 as a potent brain delivery platform for therapeutic antibodies, enabling high brain penetrance and functional efficacy in relevant CNS disease models.

### Progranulin delivery into *GRN*^-/-^ mouse brains

To further assess the versatility of SMS2 as a brain delivery platform, we evaluated its capacity to transport human progranulin (hPGRN) into the CNS. PGRN is a secreted glycoprotein essential for proper lysosomal function and brain homeostasis (*25–27*). Loss-of-function mutations in the *GRN* gene, which encodes PGRN, account for approximately 10–15% of familial FTD cases. These mutations often result in haploinsufficiency and are associated with lysosomal dysfunction, glial activation, and progressive cortical neurodegeneration. While recombinant PGRN exhibits poor BBB penetration, emerging strategies that enhance CNS delivery of PGRN have demonstrated potential to restore lysosomal function and ameliorate FTD-related phenotypes (*27–29*).

To preserve endogenous trafficking and lysosomal targeting, we fused SMS2 to the N-terminus of hPGRN, maintaining its native C-terminal lysosomal sorting motifs (Fig. 5A-B). In bEnd.3 cells, SMS2-hPGRN displayed robust mucin-dependent internalization, whereas unconjugated hPGRN exhibited minimal uptake (Fig. 5C-D). *Grn*^-/-^ mice, which recapitulate key age-dependent pathological features of GRN deficiency, including lysosomal dysfunction, neuroinflammation, and age-dependent neuronal loss (*30*), were used to evaluate *in vivo* efficacy of brain delivery. Mice were treated with saline, 15 mg/kg of SMS2-hPGRN, or an equimolar dose of unconjugated hPGRN, and bis(monoacylglycero)phosphate (BMP) levels in brain hemispheres were quantified by lipidomics (Fig. 5E). BMP depletion is an early hallmark of lipid dysregulation in FTD-GRN and serves as a sensitive readout of lysosomal health (*25, 27*). Several key genotype-dependent BMP species, including BMP(18:1/18:1) and BMP(22:6/22:6), are reduced by ∼50-70% in *Grn⁻/⁻* brains relative to wild-type as early as two months of age (*27*). While unconjugated hPGRN administration did not increase in BMP levels in *Grn*^-/-^ mice, SMS2-hPGRN treatment significantly improved BMP levels, including a 56% increase in BMP(18:1/18:1) and a 60% increase in BMP(22:6/22:6), relative to saline-treated knockout mice. These findings demonstrate that SMS2-hPGRN enhances lysosomal function and facilitates correction of GRN-deficiency-associated lipid abnormalities (Fig. 5F). More broadly, they highlight the modularity and therapeutic potential of SMS2 as a brain delivery platform, showing that it can be flexibly fused to either terminus of cargo proteins, thereby preserving critical cargo functional domains and enabling customizable strategies for CNS-targeted delivery.

**Figure 5.**
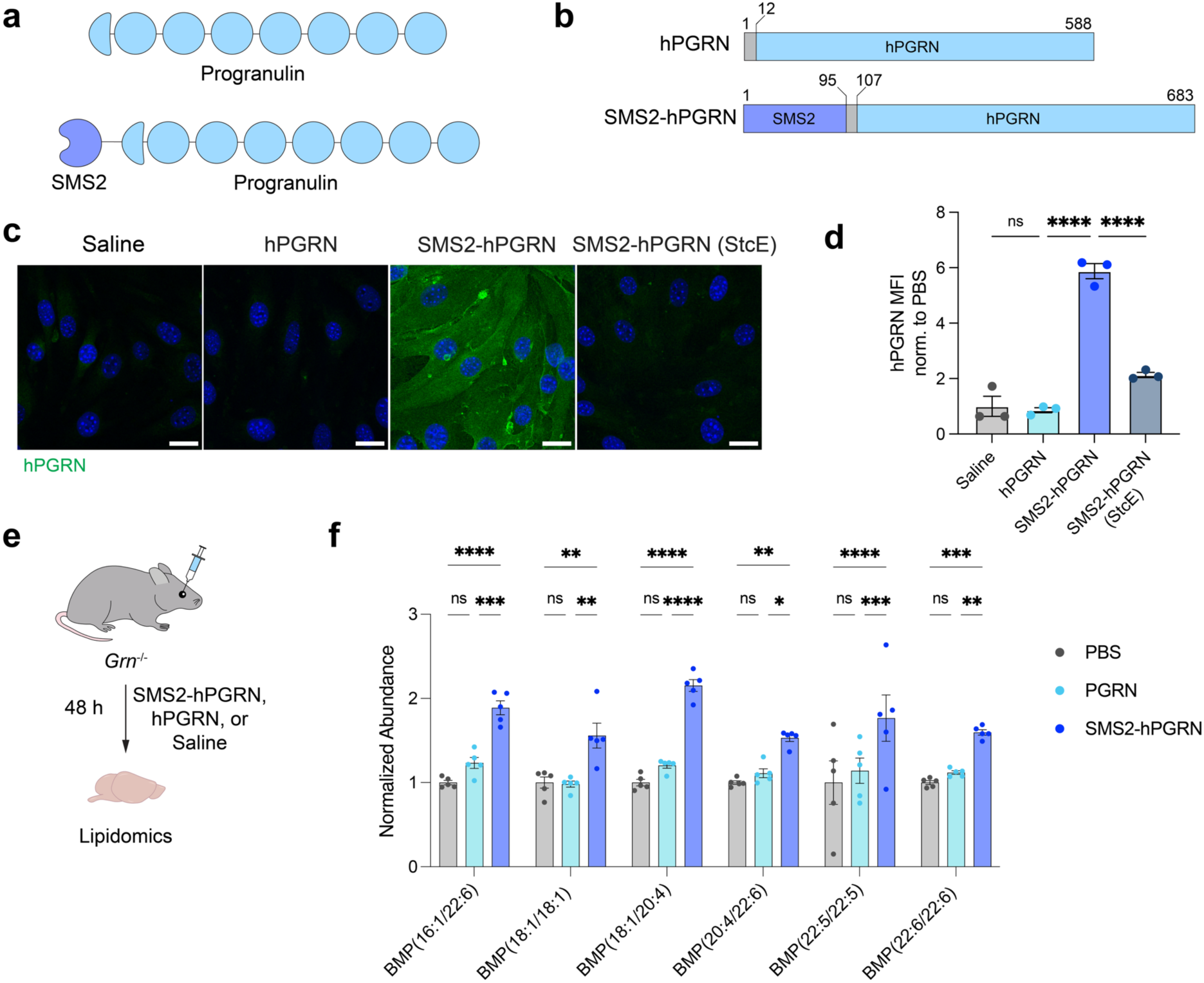
SMS2-PGRN improves rescue of lysosomal lipid dysregulation in GRN-deficient mice. a) Diagram of hPGRN and SMS2-hPGRN protein constructs. b) hPGRN and SMS2-hPGRN constructs used in this study. Highlighted regions include a GGS linker (dark gray), progranulin (blue), and SMS2 module (purple). Amino acid positions are indicated. c) Binding and internalization of hPGRN and SMS2-hPGRN in bEnd.3 cells. Binding and internalization of SMS2-hPGRN is dependent on cell surface mucins. Scale bar=25 µm. d) Quantification of hPGRN MFI in (c) (n=3 wells per construct; one-way ANOVA with Šidák’s post hoc test; mean ± s.e.m.). e) Experimental scheme for *Grn*^-/-^ mice treatment with saline, hPGRN, or SMS2-hPGRN at equimolar dosing (equivalent to 15 mg/kg SMS2-hPGRN) for 48 h. f) BMP quantification in brains of *Grn*^-/-^ mice treated with saline, hPGRN, or SMS2-hPGRN, with key altered BMP species shown (n=5 animals per group; two-way ANOVA with Tukey’s post hoc test; mean ± s.e.m.).

## Discussion

The BBB remains a central challenge in the treatment of neurological and neurodegenerative diseases, where therapeutic efficacy often hinges on achieving sufficient CNS exposure. Here, we introduce GlycoShuttles as a new glycan-targeted approach for brain delivery that leverages the brain endothelial glycocalyx as a novel entry portal across the BBB. We engineer SMS1 and SMS2 as versatile brain delivery shuttles capable of transporting diverse protein cargos into major brain cell types across brain regions. Our findings reveal that SMS-mediated internalization operates through a mucin-dependent, caveolae- and lipid raft-associated endocytic pathway, offering a mechanistically distinct alternative to RMT and other conventional BBB shuttle strategies.

We show that SMS2 preserves delivery efficiency of full-length SMS1, despite being significantly more compact (∼11 kDa vs. ∼98 kDa), which offers key translational advantages including improved biodistribution, reduced immunogenic potential, and greater compatibility with larger or structurally constrained therapeutic payloads. Notably, SMS2 exhibits favorable pharmacokinetics and achieves broad delivery across multiple brain cell types, including neurons and microglia, while reducing off-target accumulation in peripheral organs such as the liver and heart.

Using therapeutically relevant cargos, we demonstrate the versatility of SMS2 across multiple disease contexts. In a mouse model of AD, an SMS2-conjugated αBACE1 antibody achieved high levels of brain penetration, robust target engagement, and a significant reduction in brain and plasma Aβ42 levels, showing considerable promise of SMS2 for mediating antibody delivery into the brain. In a second model of GRN-FTD, we demonstrated that SMS2-fused hPGRN could be successfully delivered into the brains of *Grn* ^-/-^ mice. SMS2-hPGRN enhanced BMP levels, a sensitive biomarker of lysosomal dysfunction, more effectively than hPGRN alone, underscoring the utility of SMS2 in treating lysosomal storage disorders and protein deficiencies. The N-terminal fusion design preserved critical C-terminal lysosomal targeting motifs of hPGRN, showcasing SMS2’s modularity and compatibility with therapeutics requiring spatial or structural constraints.

Together, our findings position SMS2 as a new platform for noninvasive, systemic delivery of therapeutic cargo into the brain. By targeting the cerebrovascular glycocalyx, an abundant and underexplored feature of the brain endothelial cell surface, SMS2 unlocks a mechanistically distinct route for BBB transport that complements and expands upon RMT strategies. By engaging shared glycan motifs across multiple glycoproteins, SMS2 may circumvent limitations such as receptor saturation or pathway disruption that constrain single-receptor shuttle systems. Its compact size, efficient CNS penetrance, and modular fusion capability with diverse therapeutic cargos make SMS2 a compelling candidate for broad application across neurological diseases.

## Acknowledgments

We thank D. J. Shon, A. Chiariello, D. Selvan, and A. Hugenmatter for support with early construct studies and all members of the Wyss-Coray and Bertozzi labs for feedback and support. This work was funded by the National Institute on Aging (R01-AG072255 to T.W.-C. and RF1-AG064897 to T.W.-C.), the Phil and Penny Knight Initiative for Brain Resilience (T.W.-C., E.N.W., and M.A.R.), the National Cancer Institute (R01-CA200423 to C.R.B.), and the Alzheimer’s Association (ADSF-24-1345203-C to M.A.R.). S.M.S. was supported by a Stanford Bio-X Fellowship. G.S.T. was supported by a Hertz Foundation Fellowship, a U.S. National Science Foundation Graduate Research Fellowship, and the Stanford Sarafan Chemistry/Biology Interface Predoctoral Training Program. J.X. was supported by a Knight Initiative Postdoctoral Fellowship for Brain Resilience.

## Author contributions

S.M.S., C.R.B., and T.W.-C. conceptualized the study. G.S.T. and S.M.S. designed, purified, and validated *E. Coli*-expressed recombinant proteins. S.M.S. designed constructs for mammalian expression and purification. S.M.S., J.K.B., and H. I. P. performed and analyzed *in vitro* and *in vivo* confocal imaging experiments and conducted flow cytometry and animal experiments. J.X. performed lipidomics experiments. J.H.M. and E.N.W. performed hIgG and Aβ ELISA analysis. S.M.S. wrote the manuscript with comments from all authors. T.W.-C., C.R.B., and M. A.-R. supervised the work.

## Competing interests

T.W.-C. is a co-founder of Qinotto Inc., Teal Rise Inc., and Vero Biosceinces. C.R.B. is a co-founder and scientific advisory board member of Lycia Therapeutics, Palleon Pharmaceuticals, Enable Bioscience, Redwood Biosciences (a subsidiary of Catalent), OliLux Bio, InterVenn Bio, Firefly Bio, Neuravid Therapeutics, Valora Therapeutics, Euler Biologics, TwoStep Therapeutics, and ResNovas Therapeutics. C.R.B is also on the Board of Directors of Eli Lilly, OmniAb, Xaira Therapeutics, and Acepodia. S.M.S., T.W.-C., and C.R.B. are co-inventors on a patent application related to the work published in this paper. The remaining co-authors do not have any competing interests to declare.

## Data and materials availability

All data are available in the main text or supplementary materials. Sequences for constructs generated in this study are available upon request. Correspondence and requests for materials should be addressed to S.M.S., T.W.-C., and C.R.B.

## Materials and Methods

### Cell culture

All cells were maintained at 37°C and 5% CO_2_. bEnd.3 (ATCC, CRL-2299) and HeLa (ATCC, CCL-2) cells were cultured in high glucose DMEM (Thermo Fisher Scientific, 10567022) supplemented with 10% fetal bovine serum (FBS) and 1% penicillin/streptomycin. Human primary brain microvascular endothelial cells (Cell Systems, ACBRI 376) were cultured in Complete Classic Media with Serum (Cell Systems, 4Z0-500) according to manufacturer’s recommendations.

### Animals

C57BL/6 mice (Jackson Laboratory, stock #000664), Grn^−/−^ mice (Jackson Laboratory, stock #013175), and 5XFAD mice (Jackson Laboratories, stock #034848) were obtained from Jackson Laboratory. Mice were housed in a 12-h light/dark cycle, temperature (20-22°C) and humidity-controlled environment and provided *ad libitum* access to food and water. Both male and female mice were used, depending on availability from the vendor or in-house colony. Specifically, 3- to 4-month-old male mice were used in experiments involving mCherry constructs; 2.5-month-old female 5XFAD mice were used for anti-BACE1 antibody delivery studies; and 3-month-old male and female *Grn*⁻/⁻ mice were used in progranulin delivery studies, with data from both sexes pooled due to the absence of sex-dependent differences. All animal care and procedures complied with the Animal Welfare Act and were in accordance with institutional guidelines and approved by the institutional administrative panel of laboratory animal care at Stanford University.

### Cloning

SMS constructs were cloned from a pET28b-StcE^E447D^ plasmid using Q5 site-directed mutagenesis and HiFi DNA Assembly (New England Biolabs). The amino acid sequence for mCherry was reverse translated and optimized for expression in *E. coli* K12 with the IDT Codon Optimization Tool before cloning as described above. The anti-BACE1 antibody sequence was obtained from a previous publication (*22*). Constructs comprising PGRN and the anti-BACE1 antibody were optimized and cloned for mammalian cell expression by Genscript.

### Protein purification

ClearColi BL21(DE3) electrocompetent cells (Lucigen) cells were transformed with sequence-confirmed plasmids and grown in sterile LB-Miller broth with 30 µg/mL kanamycin at 37 °C and 250 rpm. Cultures were induced at an optical density of 0.6–0.8 with 0.4 mM isopropyl-β-D-1-thiogalactopyranoside (IPTG) and grown overnight at 20 °C and 250 rpm. Cells were centrifuged at 6,000g for 10 min and lysed with B-PER Complete Bacterial Protein Extraction Reagent (Thermo Fisher Scientific). Lysates were clarified by centrifuging at 16,000g for 20 min. Lysates were applied to 3–4 mL of Ni-NTA agarose (Qiagen) per liter of bacterial culture, washed with 200 ml of 20 mM HEPES (pH 7.5), 500 mM NaCl and 20 mM imidazole, and eluted into 1-2 ml fracitons with 20 ml of 20 mM HEPES (pH 7.5), 500 mM NaCl and 250 mM imidazole per liter of culture. Fractions that were confirmed to have eluted fractions via NanoDrop were combined and buffer exchanged into cold 2X PBS with Zeba Spin desalting columns (Thermo Fisher Scientific) or Slide-A-Lyzer dialysis cassettes (Thermo Fisher Scientific) and further purified by size exclusion chromatography using a Superdex 200 Increase 10/300 GL Column (Cytiva Life Sciences) in 2X PBS, pH 7.4. Proteins were run through Pierce high-capacity endotoxin removal columns (Thermo Fisher Scientific) at least two times, following the manufacturer’s instructions. Endotoxin levels were tested using a HEK-Blue LPS detection kit 2 (InvivoGen) according to the manufacturer’s recommendations. All endotoxin levels were confirmed to be below K/M for the maximum in vivo dose, where K is 5 EU kg^-1^ and M is the dose of the formulation within a 1-h period. Purified, endotoxin-free PGRN and anti-BACE1 constructs were produced in HEK293T cells and TurboCHO cells, respectively, at Genscript. Protein concentration was determined by NanoDrop, and protein purity was determined by SDS–PAGE (Fig. S5-7).

### Cell surface flow cytometry-based binding assays

For the StcE-treated conditions, K562 cells were cultured for 24 hours with 15 nM StcE. K562 cells (5 x 10^5^ cells) were added to each well of a V-bottom 96-well plate, washed 2X with cold fluorescence-activated cell sorting (FACS) buffer (0.5% bovine serum albumin (BSA) in PBS), once with FACS buffer containing 2 mM EDTA. Cells were treated with GlycoShuttles in cold FACS buffer and 0.1% benzonase for 30 min on ice protected from light, washed twice with FACS buffer containing 2 mM EDTA, washed once with FACS buffer, and stained with APC anti-His (clone GG11-8F3.5.1; Miltenyi Biotec) at manufacturer recommended dilution in FACS buffer for 30 min on ice protected from light. Cells were washed three times with FACS buffer containing 2 mM EDTA, stained with 5 nM Sytox Blue in FACS buffer with 2 mM EDTA for 20 min. Cells were analyzed using a NovoCyte Quanteon flow cytometer (Agilent), compensated with single stained cell controls, and analyzed using FlowJo software (BD).

### ColabFold modeling

Protein sequences for mCherry-SMS1, mCherry-SMS1ΔINS, mCherry-SMS1ΔC, and mCherry-SMS2 were used as input for ColabFold (*31*). The predicted structure for SMS1 variants aligned well with the crystal X-ray structure determined by Yu et al. (*17*) based on root mean squared deviation (RMSD) < 1.5. Molecular diagrams were generated using Pymol.

### In vitro cellular internalization assays

Cells were plated on round coverslips (EMS, 72196-12) in a 24-well plate about 16-24h before using. StcE treatment was performed using 1 nM StcE for 1h at 37°C. For endocytosis inhibitor experiments, the inhibitors were added for 20 min before co-incubation of shuttle constructs. The following final concentrations of inhibitors were used: 60U/mL nystatin (Sigma, N9150), 5mM methyl-Δ-cyclodextrin (Sigma, C4555), and 50 nM bafilomycin (Cell Signaling, 54645). For bEnd.3 targeted knockout (KO) experiments, KO cell lines were generated using Cas9 ribonucleoprotein (RNP) electroporation with guide sequences designed via Synthego. Shuttle constructs were added at 250 nM, unless otherwise noted, for 30 min and then washed 3 times with PBS before fixation in 4% paraformaldehyde for 15 min. Cells were washed 3 times with PBS for 5 minutes and blocked in 3% normal donkey serum with 0.3% Triton X-100 in PBS for 1 h at room temperature. Cells were incubated at room temperature in blocking solution with the following primary antibodies for 1.5 h: goat anti-CD31 (1:100, R&D, AF3628), rabbit anti-CAV1 (1:100, Cell Signaling Technologies, 3267S), mouse anti-CLTC (1:100, Thermo, MA1-065), rabbit anti-mCherry (1:100, Cell Signaling Technologies, 43590), rat anti-mCherry (1:100, Thermo, M11217), goat anti-RAB5 (1:100, LSBio, LS-B12415-300), mouse anti-EEA1 (1:100, Cell Signaling, 48453S), mouse anti-RAB7 (1:100, Cell Signaling, 95746S), rabbit anti-RAB11 (1:100, Cell Signaling, 5589S), goat anti-golgi protein 58k (1:100, Thermo PA1-9000), rat anti-LAMP1 (1:100, Thermo, 14-1071-82), and goat anti-progranulin (1:100, Biotechne, AF2420). Cells were subsequently washed three times with PBS, stained with the appropriate Alexa Fluor-conjugated secondary antibodies (1:500, Thermo) for 1 h at room temperature, washed three times again, mounted, and coverslipped with Vectashield Hardset Antifade Mounting Medium with DAPI (Vector Labs, H-1500-10) or ProLong Gold Antifade Mountant (Thermo Fisher Scientific, P36934). Imaging was performed on a confocal laser-scanning microscope (Zeiss LSM880), and images were analyzed using ImageJ.

### In vivo immunofluorescence analysis

Mice were injected with retro-orbitally with constructs at the specified doses. Mice were anesthetized with Avertin and perfused with PBS before tissue extraction. Tissues were fixed in 4% PFA at 4°C overnight and preserved in 30% sucrose in PBS for at least 24 h before sectioning. Tissues were sectioned into 40 μm slices using a microtome (Leica), blocked in 3% normal donkey serum with 0.3% Triton X-100 in TBS-T for 1.5 h at room temperature, and then incubated at 4°C overnight with the following primary antibodies: goat anti-CD31 (1:100, R&D, AF3628), goat anti-Iba1 (1:100, Abcam, ab5076), mouse anti-NeuN (1:200, Sigma, MAB377), chicken anti-GFAP (1:1000, Thermo, PA1-10004), rabbit anti-mCherry (1:100, Cell Signaling Technologies, 43590), and rat anti-mCherry (1:100, Thermo, M11217). The following day, slices were washed three times with TBS-T, stained with the appropriate Alexa Fluor-conjugated secondary antibodies (1:250, Thermo Fisher Scientific) for 2 h at room temperature, washed three times again, mounted, and coverslipped with Vectashield Hardset Antifade Mounting Medium with DAPI (Vector Labs, H-1500-10). Imaging was performed on a confocal laser-scanning microscope (Zeiss LSM880), and images were analyzed using ImageJ and Imaris 10.

### Flow cytometry quantification of mCherry in brains

Following anesthetization of mice with Avertin and PBS perfusion, brains were extracted and prepared for flow cytometry as previously described (*32*). Briefly, hemibrains were minced and digested with a neural dissociation kit (Miltenyi, 130-092-628) according to manufacturer’s instructions. Brain homogenates were filtered through a 100 μm strainer and then centrifuged in 0.9 M sucrose to remove myelin. Samples were blocked for 10 min with Fc preblock (CD16/CD32, BD, 553141) and stained for 30 min with rat anti-CD31-FITC (1:100, Biolegend, 102406), rat anti-CD45-PE/Cy7 (1:200, Biolegend, 103114), rat anti-CD90-FITC (1:100, Biolegend, 105305), and rat anti-CD11b-APC (1:100, Biolegend, 101212). Live cells were identified using Sytox Blue viability dye (1:1000, Thermo Fisher Scientific, S34857). Flow cytometry analysis was performed on a Sony SH800S sorter, and data were analyzed using FlowJo software (TreeStar). mCherry signal was quantified in live, singlet cells for each sample with gating of brain endothelial cells (CD31^+^), neurons (CD90^+^), and microglia (CD11b^+^) when applicable.

### Vascular depletion and validation

Brain samples were homogenized in HBSS using a Dounce homogenizer as previously described (*33*). Homogenized tissue was centrifuged at 1000g for 10 min at 4°C to pellet microvessels and cells. The supernatant was saved as the non-cellular associated fraction on ice. The cell pellet was mixed with 25% dextran in HBSS and centrifuged at 4400g for 20 min at 4°C. The pellet was saved as vascular cells on ice. The supernatant was transferred to a new tube, diluted with 20mL of HBSS and centrifuged at 4400g for 20 min at 4°C. The pellet was saved as parenchymal cells on ice. Cell fractions were lysed in 1X RIPA lysis buffer (Thermo) supplemented with 1x cOmplete™ protease inhibitor cocktail (Sigma) and centrifuged at 13,000g for 15 min at 4°C. Protein concentrations were measured by BCA (Pierce) and normalized before quantification of mouse CD31 via ELISA according to manufacturer instructions (Abcam, ab204527).

### Tissue ELISA analysis

Tissues were homogenized via sonication in 1X RIPA lysis buffer (Thermo) supplemented with 1x cOmplete™ protease inhibitor cocktail (Sigma). Lysates were centrifuged at 13,000g for 15 min at 4°C, and supernatant protein concentrations were measured by BCA (Pierce). mCherry ELISA quantification (Abcam, ab221829) was performed on mouse brain tissue and plasma according to manufacturer instructions. Human Aβ42 and Aβ40 were measured in mouse plasma and cortex samples using the V-PLEX Aβ Peptide Panel 1 (6E10; Cat# K15200E, Meso Scale Diagnostics, Rockville, MD) in a blinded manner. Assay was run according to manufacturer directions and plates were measured using the MESO QuickPlex SQ 120 instrument (Meso Scale Diagnostics). Human immunoglobulin isotyping panel 1 kit (Cat# K15203D, Meso Scale Diagnostics) was used to measure human Ig concentration in mouse cortex and blood samples in a blinded manner. Assays were run according to manufacturer directions and plates were measured using the MESO QuickPlex SQ 120 instrument (Meso Scale Diagnostics). To demonstrate specificity of the human IgG assay, human isotypes IgA and IgM were also measured and were not detected.

### Plasma PK/PD

For plasma PK/PD evaluation, mice were dosed intravenously with the specified construct and in-life plasma was take via saphenous vein bleeding. Blood was collected in EDTA- or heparin-coated tubes and centrifuged at 2000g for 15 min at 4°C. Plasma was isolated for subsequent analysis via ELISA assays as described above.

### Targeted BMP lipidomics

Following PBS perfusion of mice, brains were extracted, snap-frozen on dry ice, and stored at −80 °C until further processing as previously described (*28, 34*). Briefly, frozen left hemisphere brain tissue was homogenized in a dounce tissue homogenizer in 1 mL of potassium phosphate buffered saline (KPBS, 136 mM KCl, 10 mM KH_2_PO_4_, pH 7.4). For lipid extraction, 25 µL of homogenate were added to a microcentrifuge tube containing 1 mL of chloroform:methanol (2:1, v/v) containing SPLASH LIPIDOMIX (750 ng/mL, Avanti) and vortexed for 1 h at 4 °C. Subsequently, 200 µL of 0.9% (w/v) NaCl solution (VWR) was added, and the mixture was vortexed again for 10 min at 4 °C and centrifuged at 3000 x g for 5 min. After centrifugation, the mixture was separated into aqueous layer (upper) and organic layer (bottom). 600 µl of organic layer were transferred into a new microcentrifuge tube and dried in speedvac until all solvent was evaporated. Dried lipids were then solubilized in 50 µl of lipid reconstitution buffer (acetonitrile: isopropanol: H_2_O 13:6:1, v/v/v) and vortexed for 10 minutes at 4 °C. The samples were centrifuged at 20,627g for 10 min at 4 °C, and 45 µl was transferred into a glass vial with inserts for mass spectrometry analysis. Lipids were separated on an Agilent RRHD Eclipse Plus C18, 2.1 × 100 mm, 1.8u-BC column with an Agilent guard holder (UHPLC Grd; Ecl. Plus C18; 2.1 mm; 1.8 µm). Before mass spectrometry, the columns were connected to a 1290 LC system for lipid separation. The liquid chromatography system was linked to an 6495D Triple Quadrupole (QQQ) mass spectrometer with a liquid chromatography–electrospray ionization probe. External mass calibration was performed weekly using a QQQ standard tuning mix. The column compressor and autosampler were held at 45 °C and 4 °C, respectively. The mass spectrometer parameters included a capillary voltage of 4.4 kV in positive mode and 5.5 kV in negative mode, and the gas temperature and sheath gas flow were held at 200 °C and 275 °C, respectively. The gas flow and sheath gas flow were 10 and 11 L per min, respectively, whereas the nebulizer was maintained at 45 psi. The nozzle voltages were maintained at 500V in the positive mode and 1,000V in the negative mode. 3 µL injection volumes were used for each sample for polarity switching. A mobile phase with two distinct components (A and B) was used in the chromatographic process. Mobile phase A was a mixture of acetonitrile:water (2:3 (v/v)), whereas mobile phase B was composed of isopropanol:acetonitrile (9:1 (v/v)), both containing 0.1% formic acid and 10 mM ammonium formate. The elution gradient was carried out over a total of 16 min, with an isocratic elution of 15% B for the first minute, followed by a gradual increase to 70% of B over 3 min and then to 100% of B from 3 to 14 min. Subsequently, this was maintained from 14 to 15 min, after which solvent B was reduced to 15% and maintained for 1 min, followed by an extra 2 min for column re-equilibration. The flow rate was set to 0.400 mL/min. The QQQ was set to operate in multiple reaction monitoring (MRM) to analyze compounds of interest. Standard lipids, glycerophosphodiesters and amino acids were optimized using the MassHunter Optimizer MRM, a software used for automated method development. For most species, the two most abundant transitions were selected to detect it. The precursor–product ion pairs (*m*/*z*) of the compounds used for MRM are listed in Supplementary Table 1. Annotation and quantification of lipids were performed using Quant-My-Way (Agilent). Individual lipid species shown in the figures were validated using the Qualitative software by manually checking the peak alignment and matching the retention times and MS/MS spectra to the characteristic fragmentation compared to the standard compounds. Analyzing two transitions for the same compound and looking for similar relative response was an added validation criterion to ensure the correct species were identified and quantified. The MRM method and retention time were used to quantify all lipid species using the quantification software, and the raw peak areas of all species were exported to Microsoft Excel for further analysis. Raw abundances were normalized to PC (18:1/18:1).

## Supplementary Figures

**Fig. S1.**
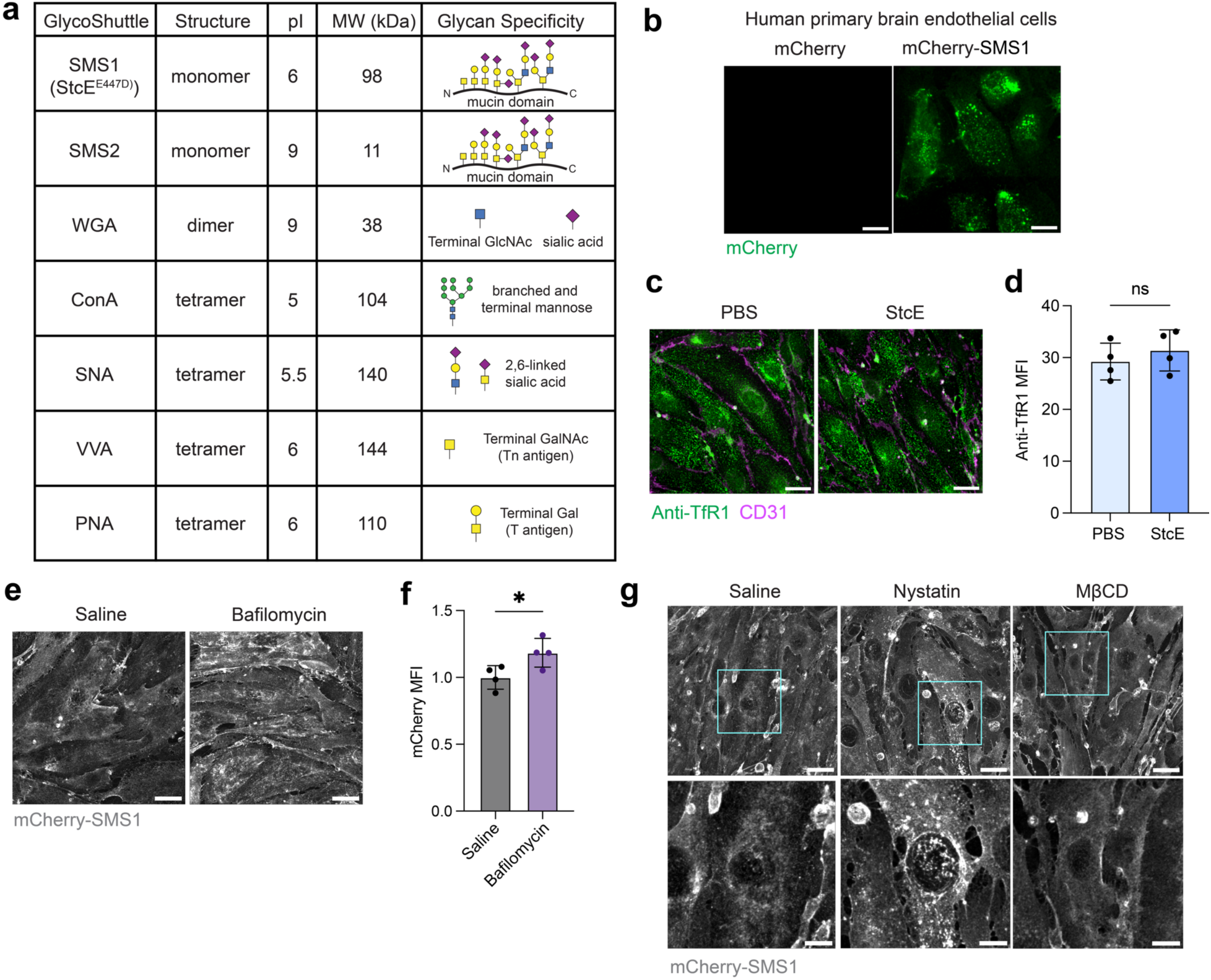
Characterization of GlycoShuttle constructs. a) Properties of glycocalyx-binding proteins tested including quaternary structure, molecular weight (MW), isoelectric point (pI), and glycan specificity (*9*). b) SMS1 shuttles mCherry cargo into human primary brain endothelial cells. Scale bar=25 µm. c) Anti-TfR1 (8D3) internalization into bEnd.3 cells is unaffected by StcE-mediated mucin removal. Scale bar=25 µm. d) Quantification of anti-TfR1 MFI in (c) (n=4 wells per construct; two-sided t-test; mean ± s.e.m.). e) Treatment of bEnd.3 cells with bafilomycin increases mCherry-SMS1 punctate signal. Scale bar=25 µm. f) Quantification of mCherry MFI in (f) (n=4 wells per construct; two-sided t-test; mean ± s.e.m.). g) Treatment of bEnd.3 cells with caveolae/lipid raft inhibitors perturbs mCherry-SMS1 binding and internalization into cells. Scale bar=25 µm.

**Fig. S2.**
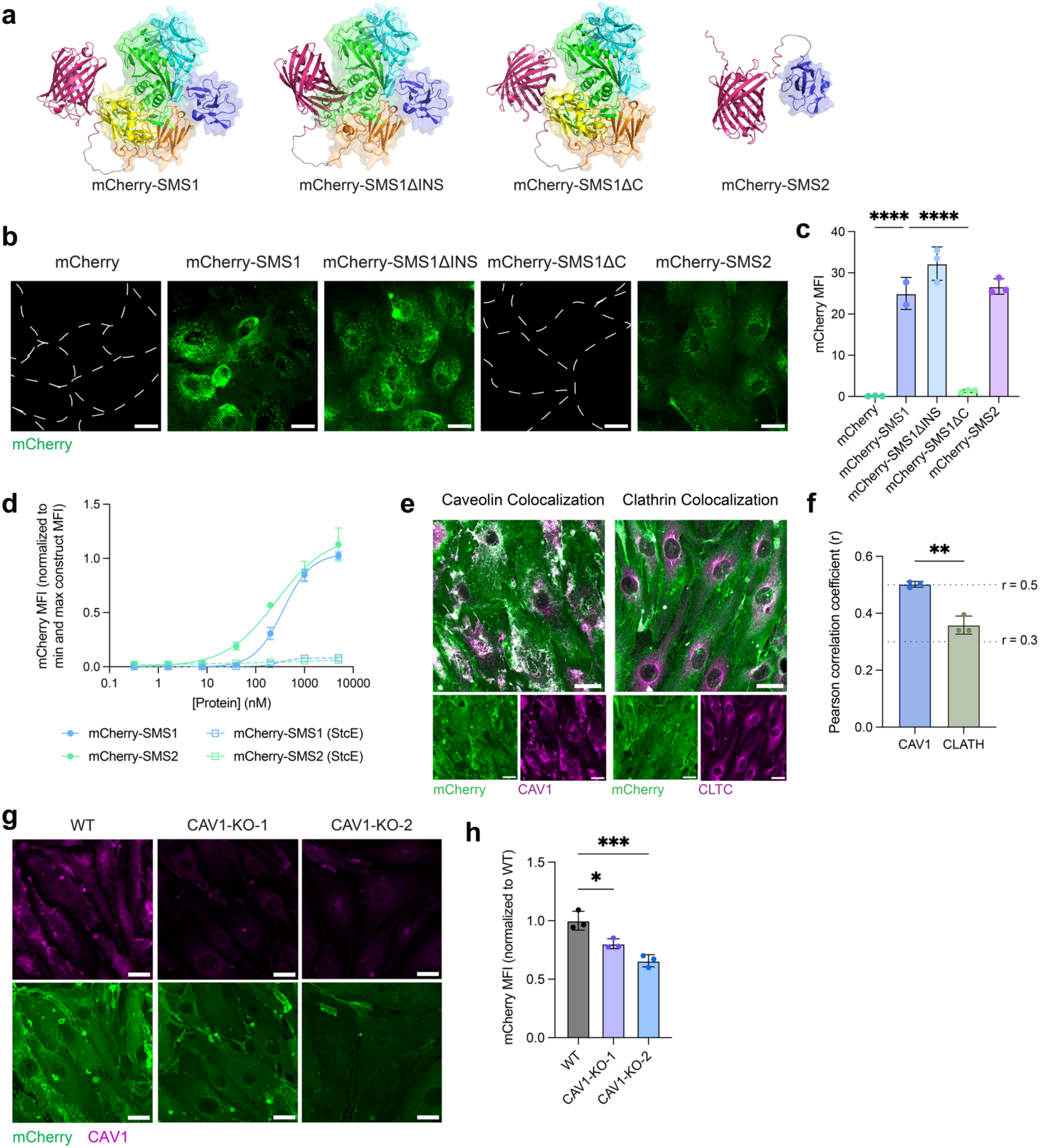
Characterization of mCherry-SMS variants. a) Structures of mCherry-SMS variants, as predicted by AlphaFold. b) Binding and internalization of mCherry-SMS1, mCherry-SMS1ΔINS, mCherry-SMS1ΔC, mCherry-SMS2, and mCherry in human primary brain endothelial cells after 30-min incubation at 37°C. Cell boundaries for mCherry-SMS1ΔC and mCherry are marked in white dotted line. Scale bar=25 µm. c) Quantification of (b) (n=2-3 wells per construct; one-way ANOVA with Dunnett’s post hoc test; mean ± s.e.m.). d) Binding curves of mCherry-SMS1 and mCherry-SMS2 on HeLa cells with and without StcE treatment. e) mCherry-SMS2 colocalization with CAV1 and CLTC in bEnd.3 cells with colocalization threshold mask in grayscale. Scale bar=25 µm. f) Colocalization analysis using Pearson correlation between mCherry-SMS2 and CAV1 or CLTC in bEnd.3 cells (n=3 wells per construct; two-sided t-test; mean ± s.e.m.). g) mCherry-SMS2 internalization in wild-type (WT) and two CAV1-KO cell lines. Scale bar=25 µm. h) Quantification of mCherry MFI in (g) (n=3 wells per construct; one-way ANOVA with Dunnett’s post hoc test; mean ± s.e.m.).

**Fig. S3.**
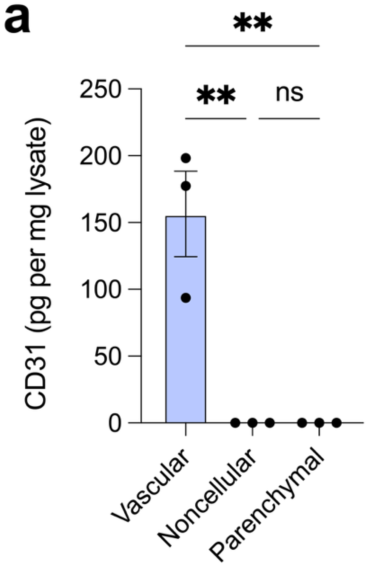
Validation of vascular depletion. a) CD31 measurement in vascular cell, non-cellular, and parenchymal cell fractions by ELISA after vascular depletion (n=3 animals; one-way ANOVA with Tukey’s post hoc test; mean ± s.e.m.).

**Fig. S4.**
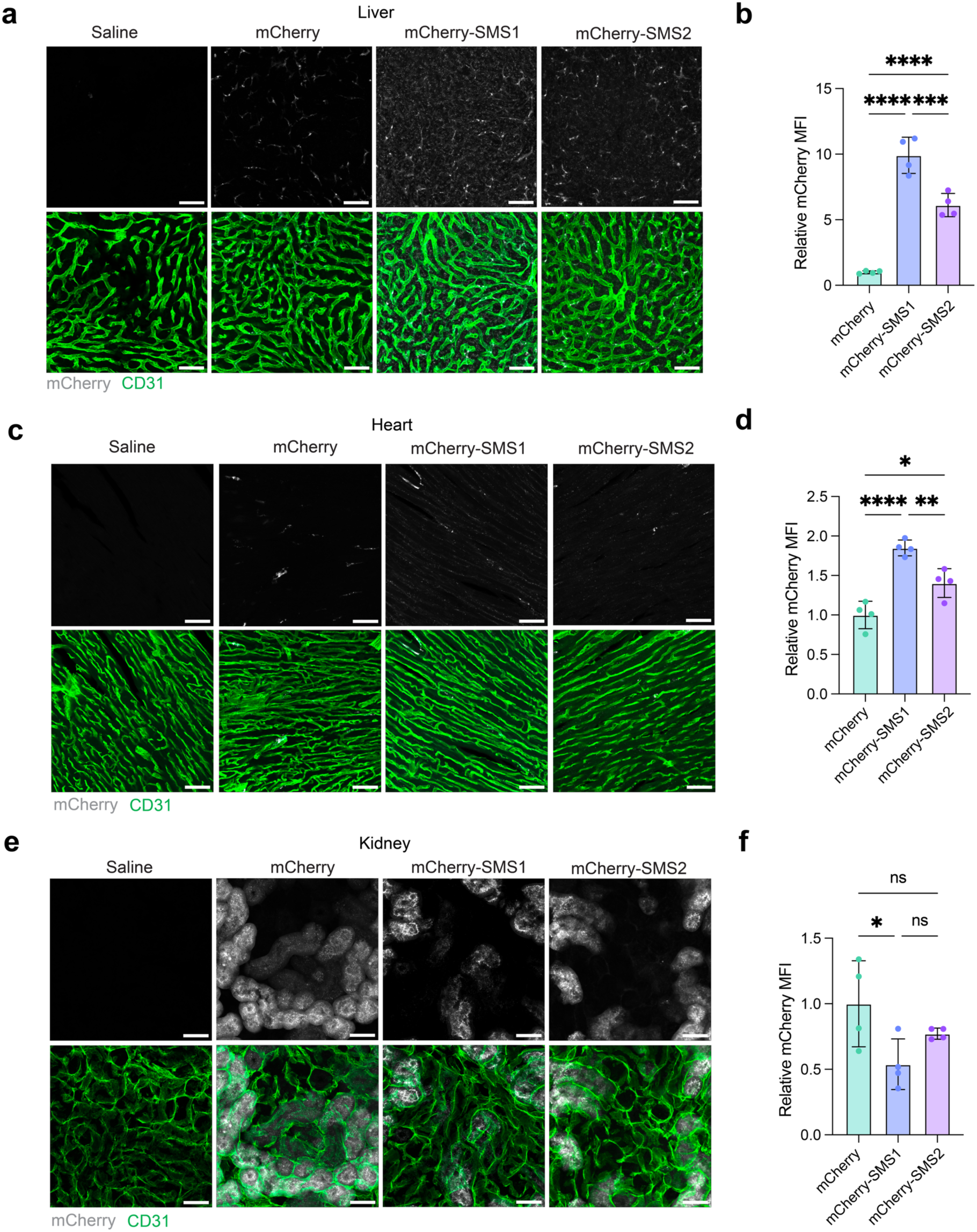
Distribution of mCherry-SMS1 and mCherry-SMS2 in peripheral organs. a) Representative liver sections showing distribution of mCherry, mCherry-SMS1, and mCherry-SMS2 at 48 hours following equimolar dosing (equivalent to 5 mg/kg mCherry-SMS1). Scale bar=50 µm. b) Quantification of relative mCherry MFI in (a) (n=4 animals per construct; one-way ANOVA with Tukey’s post hoc test; mean ± s.e.m.). c) Representative heart sections showing distribution of mCherry, mCherry-SMS1, and mCherry-SMS2 at 48 hours following equimolar dosing (equivalent to 5 mg/kg mCherry-SMS1). Scale bar=50 µm. d) Quantification of relative mCherry MFI in (c) (n=4 animals per construct; one-way ANOVA with Tukey’s post hoc test; mean ± s.e.m.). e) Representative kidney sections showing distribution of mCherry, mCherry-SMS1, and mCherry-SMS2 at 48 hours following equimolar dosing (equivalent to 5 mg/kg mCherry-SMS1). Scale bar=50 µm. f) Quantification of relative mCherry MFI in (e) (n=4 animals per construct; one-way ANOVA with Tukey’s post hoc test; mean ± s.e.m.).

**Fig. S5.**
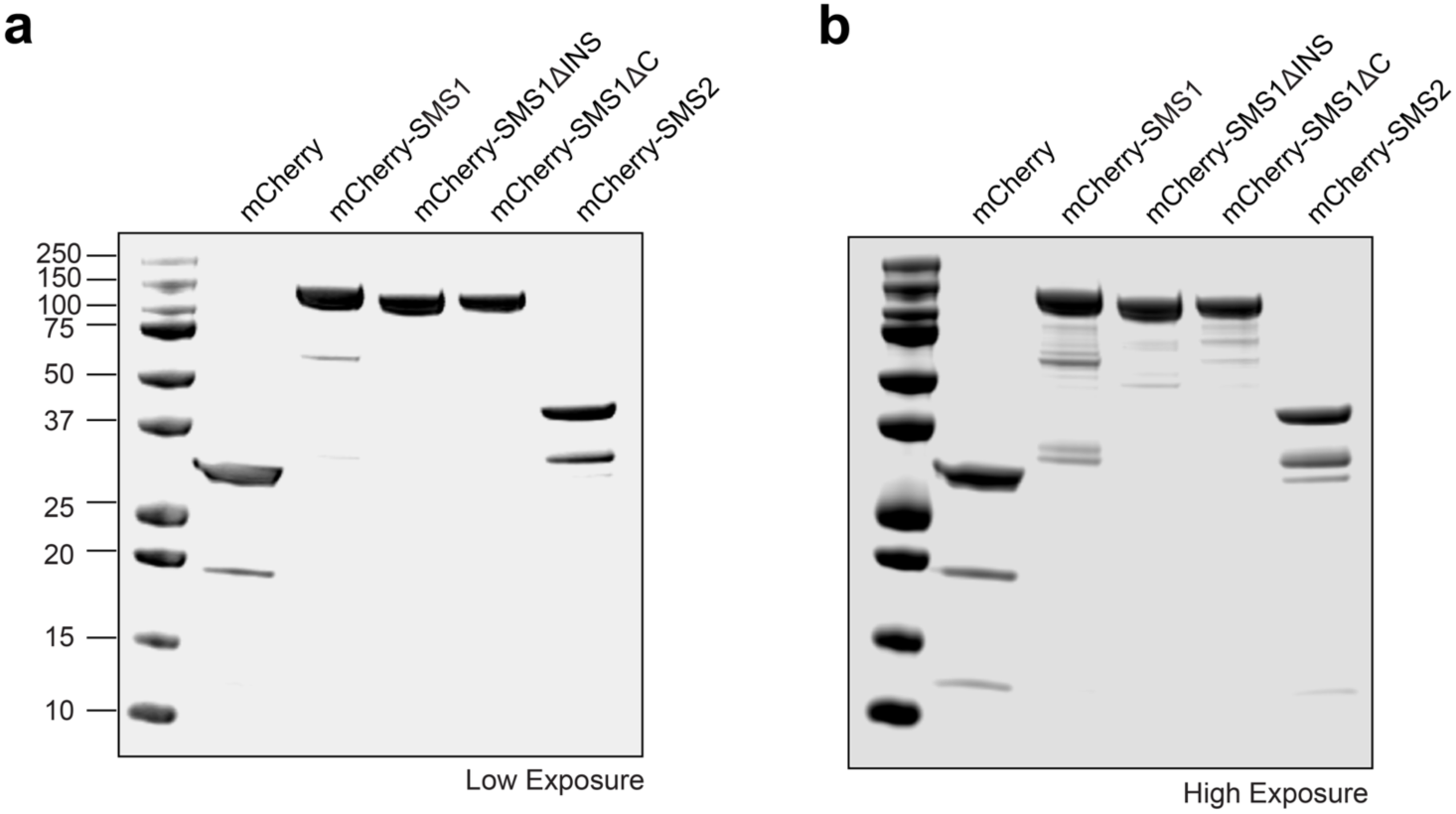
Validation of mCherry-SMS variants. a) SDS-PAGE gel of mCherry-SMS variants with low exposure. b) SDS-PAGE gel of mCherry-SMS variants with high exposure.

**Fig. S6.**
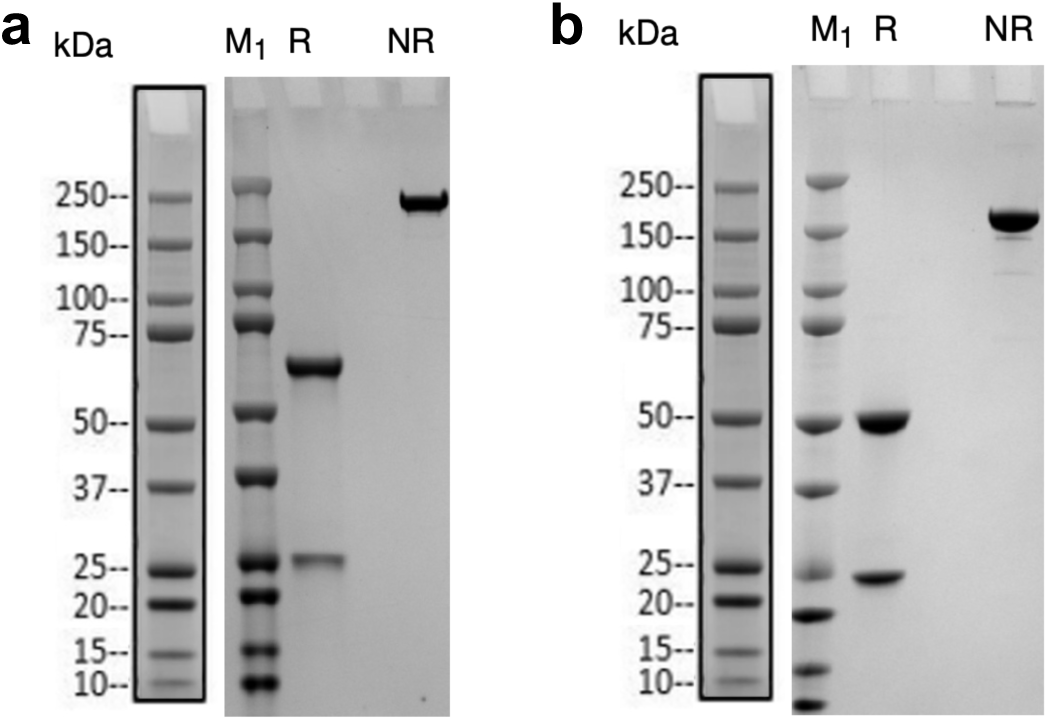
Validation of anti-BACE1 antibody constructs. a) SDS-PAGE gel of anti-BACE1-SMS2 from Genscript. R: Reducing condition; NR: Non-reducing condition; M_1_: Protein ladder. b) SDS-PAGE gel of anti-BACE1 from Genscript. R: Reducing condition; NR: Non-reducing condition; M_1_: Protein ladder.

**Fig. S7.**
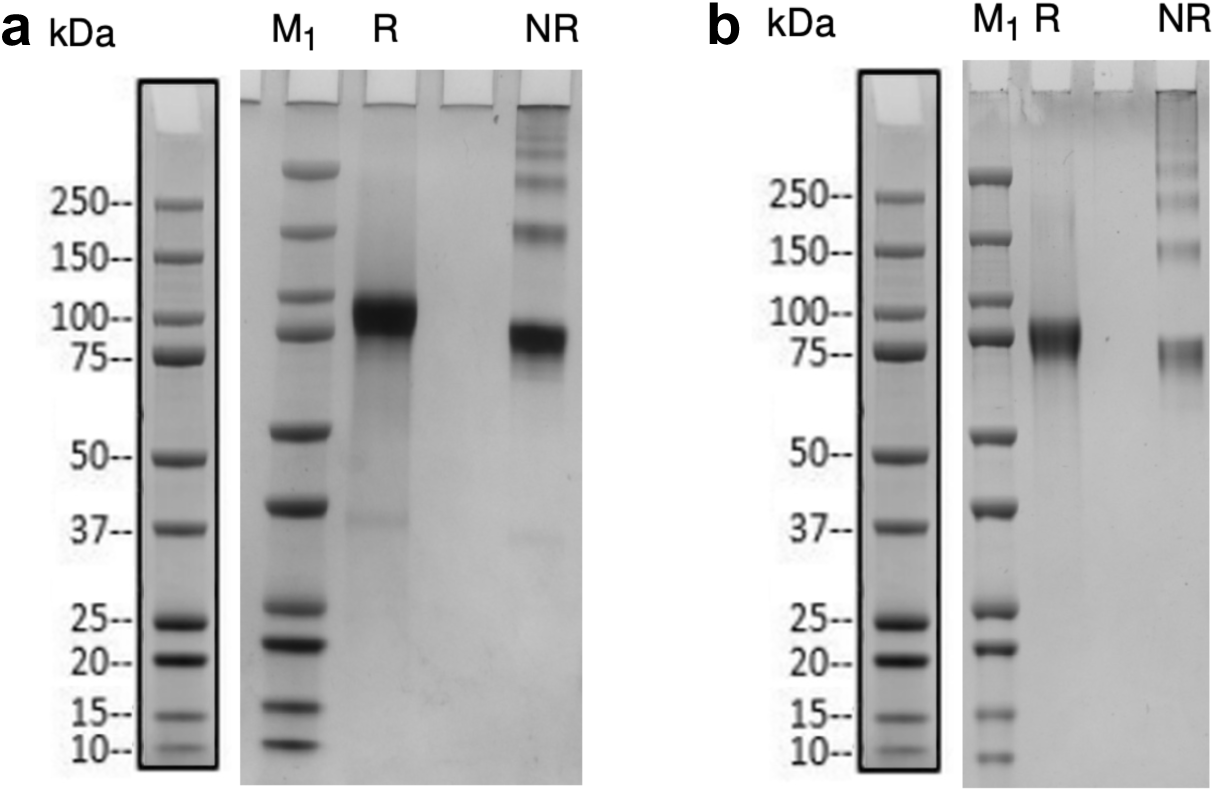
Validation of progranulin constructs. a) SDS-PAGE gel of SMS2-hPGRN from Genscript. R: Reducing condition; NR: Non-reducing condition; M_1_: Protein ladder. b) SDS-PAGE gel of hPGRN from Genscript. R: Reducing condition; NR: Non-reducing condition; M_1_: Protein ladder.

## Notes

### Summary of Updates

A misspelling in the corresponding email has been corrected.

